# NFYA promotes the anti-tumor effects of gluconeogenesis in hepatocellular carcinoma through the regulation of PCK1 expression

**DOI:** 10.1101/2022.06.30.498215

**Authors:** Goki Tsujimoto, Rin Ito, Kei Yoshikawa, Chihiro Ueki, Nobuhiro Okada

**Affiliations:** Graduate School of Interdisciplinary Science & Engineering in Health Systems, Okayama University, Okayama, Japan

**Keywords:** Hepatocellular carcinoma (HCC), Cancer metabolism, Gluconeogenesis, Cell death, Oxidative stress, NFYA, PCK1

## Abstract

Reprogramming of glucose metabolism occurs in many human tumor types, and one of these, gluconeogenesis, is known to exhibit anti-tumor effects in hepatocellular carcinoma (HCC). The transcription factor NFYA regulates gluconeogenesis in the normal liver tissue, but the function of the NFYA-gluconeogenesis axis in cancer and the functional differences of NFYA splicing variants in the regulation of gluconeogenesis is still unclear. Here, we demonstrate that NFYAv2, the short-form variant, upregulates the transcription of a gluconeogenic enzyme PCK1. We further reveal that its regulation induces high ROS levels and energy crisis in HCC and promotes cell death. These indicate that the NFYAv2-gluconeogenesis axis has enhanced anti-tumor effects in HCC, suggesting that the axis may be a potential therapeutic target for HCC. Furthermore, Nfyav1-deficient mice, spontaneously overexpressing Nfyav2, had no increasing gluconeogenesis in the liver. Taken together, our results reveal NFYAv2-gluconeogenesis axis has anti-tumor effects and the potential for NFYAv2 to be a safer therapeutic target for HCC.

## Introduction

Hepatocellular carcinoma (HCC) is the fourth leading cause of cancer-related death; moreover, it is a highly lethal tumor with a 5-year survival rate of less than 20 % for HCC patients (Villanueva, 2019; Brar et al., 2020). Therefore, developing new therapies and identifying molecular targets for treatment are essential. Recent advances in cancer metabolism research have revealed that reprogramming glucose metabolism is one of the hallmarks of cancer (Hanahan and Weinberg, 2011; Pavlova and Thompson, 2016). In particular, enhanced aerobic glycolysis, known as the Warburg effect, has been well studied in cancers, including HCC, to provide glycolytic intermediates essential for tumor cell growth (Hay, 2016). However, gluconeogenesis, a metabolic pathway that is an inverse reaction to glycolysis, has little attention. Recent studies have reported that the expression of the gluconeogenic enzymes phosphoenolpyruvate carboxykinase 1/2 (PCK1/2) and fructose-1,6-bisphosphatase 1 (FBP1) is suppressed in HCC. Furthermore, their forced expression in HCC has been shown to exert the anti-tumor effects by converting oxaloacetic acid (OAA) to phosphoenolpyruvate (PEP), which removes intermediates of the tricarboxylic acid (TCA) cycle and induces TCA cataplerosis in the cells (Ma et al., 2013; Liu et al., 2018b, 2018a; Tang et al., 2018; Grasmann et al., 2019; Wang and Dong, 2019). In addition, tumor tissues have extremely low glucose concentrations compared to normal tissues, suggesting they are suitable for the induction of gluconeogenesis (Hirayama et al., 2009). Therefore, the gluconeogenesis pathway is considered a potential therapeutic target for HCC. However, it has been suggested that the anti-tumor effects of gluconeogenic enzymes are highly context-dependent. In particular, in tumors derived from non-gluconeogenic organs, gluconeogenic enzymes have been shown to have tumor-promoting effects (Grasmann et al., 2019; Wang and Dong, 2019). Furthermore, elevated expression of gluconeogenic enzymes and increased gluconeogenesis have been reported to occur in the liver of patients with type 2 diabetes, raising concerns about its safety as a therapeutic target for HCC (Yoon et al., 2001; Barthel and Schmoll, 2003; Sakai et al., 2012). These findings create issues for therapeutic development targeting gluconeogenesis for HCC. Therefore, it is crucial to conduct a more detailed analysis of the anti-tumor effects of gluconeogenesis and to identify more selective and effective therapeutic targets.

Nuclear transcription factor Y (NF-Y) functions as a trimeric protein complex composed of three subunits, NFYA, NFYB, and NFYC, and activates the transcription of target genes with the CCAAT box on the promoter region (Dolfini et al., 2012; Gurtner et al., 2017). NFYA has a DNA-binding domain at the C-terminus and is thought to function as a limiting subunit of the NF-Y complex because point mutations in the DNA-binding domain inhibit NF-Y-DNA complex formation (Mantovanis et al., 1994). NFYA has two types of splicing variants: a long-form (NFYAv1) and a short-form (NFYAv2) that lacks the 29 amino acids encoded by exon 3, including the Q-rich transactivation domain, and these two variants have been reported to perform different functions in breast cancer (Li et al., 1992; Okada et al., 2022). In breast cancer, it has been reported that NFYAv1 regulates lipid metabolism and promotes tumor malignant behavior, while NFYAv2 has no effects. Furthermore, NF-Y has been reported to regulate gluconeogenesis in the liver and is considered one of the therapeutic targets for type 2 diabetes (Zhang et al., 2018). However, it remains unclear whether NF-Y regulates gluconeogenesis in HCC as in normal hepatocytes and whether there are functional differences in NFYA splicing variants in this process.

In this study, we investigated the functional importance of NFYA splicing variants for the anti-tumor effects of gluconeogenesis in HCC and found that the expression of NFYA, especially NFYAv2, is upregulated in HCC in response to glucose deprivation. Furthermore, we found that NFYAv2 induces high reactive oxygen species (ROS) levels and energy crisis, thereby causing cell death in glucose-deprived HCC by promoting the gluconeogenic enzyme PCK1 transcription.

## Materials and Methods

### Animal experiments

All mouse experiments were performed following Okayama University Institutional Animal Care and Use Committee-approved protocol (OKU-2019325). To analyze protein and mRNA expression levels and blood glucose concentrations, C57BL/6N mice fasted for 24 hours and then restarted feeding for 24 hours. Blood glucose levels were determined using blood from the tail and measured by a blood glucose meter for laboratory animals (ForaCare Japan, SUGL-001).

### qRT-PCR analysis

According to the manufacturer’s instruction, total RNA was extracted from the mouse liver using TRIzol reagent (Thermo Fisher Scientific, 15596018). For qRT-PCR analysis, cDNA was synthesized from total RNA using a High-Capacity RNA-to-cDNA Kit (Applied Biosystems, 4387406). mRNA expression levels were determined by qRT-PCR with KAPA SYBR FAST qPCR Master Mix Kit (Kapa Biosystems, KK4610). cDNA was synthesized from human cells using SuperPrep II Cell Lysis & RT Kit (TOYOBO, SCQ-401). mRNA expression levels were determined by qRT-PCR with KOD SYBR qPCR Mix (TOYOBO, QKD-201). Relative expression levels were normalized to mouse *Actb* or human *ACTB*. The primers used here are shown in Supplementary Table S1.

### Western blot analysis

Total cellular extracts resolved by SDS-PAGE were transferred to PVDF membranes. Western blot was performed in TBST (100 mM Tris-HCl at pH7.5, 150 mM NaCl, 0.05 % Tween-20) containing 5 % Blocking One (nacalai tesque, 03953-95). Immunoreactive protein bands were visualized using SuperSignal West Pico PLUS Chemiluminescent Substrate (Thermo Scientific, 34577).

### Cell culture

SK-Hep1, HepG2, Hep3B, Huh-4, and Huh-7 cells were maintained in DMEM supplemented with 10 % FBS and 1 % penicillin and streptomycin.

For glucose deprivation, cells were washed with PBS (-) twice and cultured in the glucose deprivation medium (DMEM (No Glucose) (Wako, 042-32255) supplemented with 2 mM sodium pyruvate (Wako, 190-14481), 10 mM sodium lactate (Sigma, L4263), and 2 mM L-glutamine (Wako, 073-05391)) for the indicated time.

### Cell proliferation

For direct cell counting experiments, SK-Hep1 cells were plated in triplicate at 0.4 × 10^4^ cells per well of a 24-well plate. At indicated days, cells were trypsinized and counted.

### CRISPR/Cas9 system

CRISPR/Cas9 system knocked out the NFYA gene in SK-Hep1 cells. NFYA and NFYAv1-specific gRNAs were determined using the candidates provided by GPP sgRNA Designer (https://portals.broadinstitute.org/gpp/public/analysis-tools/sgrna-design) and cloned into the px459 vector (Addgene, 48139). Sequence for human NFYA sgRNA #1: GCCTTACCAGACAATTAACC, human NFYA sgRNA#2: GAGCAGATTGTTGTCCAGGC, human NFYAv1 sgRNA: GCCCAGGTGGCATCCGCCTC.

### Cell survival assay

Cells were cultured in the glucose deprivation medium for 10 hours and then stained using a Double staining kit (Dojindo, 341-07381). Briefly, 10 hours later, cells were stained with 0.67 µM of Calcein-AM and 1.3 µM of PI in PBS (-) for 20 min at 37 ºC. After staining, cell images were visualized with a Keyence microscope (BZ-9000) and counted more than 600 cells to determine the proportion of cell death.

### ROS assay

Cells were cultured in the glucose deprivation medium with or without NAC and dimethyl α-KG for 3.5 hours and evaluated ROS levels using the Highly Sensitive DCFH-DA ROS Assay Kit (Dojindo, 340-09811) according to the manufacturer’s instruction. Briefly, 3.5 hours later, cells were incubated with highly sensitive DCFH-DA dye working solution at 37 ºC, 5 % CO_2_ for 30 min. After incubation, cell images were visualized with a Keyence microscope (BZ-9000) and measured fluorescence intensity in 100 cells to determine ROS levels.

### Intracellular metabolites and ATP measurements

All intracellular metabolites and ATP were measured at 5 hours after glucose deprivation using Glutamine Assay Kit-WST (Dojindo, 348-09611), Glutamate Assay Kit-WST (Dojindo, 345-09621), α-Ketoglutarate Assay Kit (Sigma, MAK054), and ATP Assay Kit (Abcam, ab83355) according to the manufacturer’s instruction. Briefly, 5 hours later, 1 × 10^6^ cells were collected and lysed, and the lysates were deproteinized with a Nanosep Centrifugal Filtration Device (10K) (Pall, OD010C33). The lysates were incubated with a working solution at 37 ºC for 30 min for intracellular metabolites or at room temperature for 30 min for ATP. After incubation, the absorbance was measured at 450 nm for glutamine and glutamate or 570 nm for α-KG and ATP.

### CUT&RUN analysis

CUT&RUN experiments were performed as previously described (Okada et al., 2022) with the CUT&RUN assay kit (CST, 86652). In brief, cells were collected 5 hours after glucose deprivation treatment, washed, bound to activated Concanavalin A magnetic beads, and permeabilized with antibody binding buffer containing Digitonin. The bead-cell complex was incubated with 0.7 µg of NFYA monoclonal antibody overnight at 4 ºC. The bead-cell complex was washed with Digitonin buffer and incubated pAG-MNase solution for 1 hour at 4 ºC. After washing with Digitonin buffer, 2 mM calcium chloride was added to activate pAG-MNase and incubated for 30 min at 4 ºC. After incubation, the reaction was stopped with a stop buffer. DNA fragments were released by incubation for 10 min at 37 ºC and purified with a Fast Gene Gel/PCR extraction kit (Nippon genetics, FG-91202). The DNA fragments were quantified by qRT-PCR analysis. The primers used here are shown in Supplemental Table S1.

### Pyruvate tolerance test (PTT)

For the pyruvate tolerance test, 16h-fasted mice were intraperitoneally injected with pyruvate (2 g/kg of body weight). Blood glucose levels were determined using blood from the tail and measured by a blood glucose meter for laboratory animals (ForeCare Japan, SUGL-001) at the indicated time.

### Antibodies

The following monoclonal (mAb) and polyclonal (pAb) primary antibodies were used for western blot: NFYA pAb (1:500; Santa Cruz, sc-10779), NFYA mAb (1:500; Santa Cruz, sc-17753), PCK1 mAb (1:1000; Santa Cruz, sc-271029), G6PC pAb (1:1000; Proteintech, 22169-1-AP), E-Cadherin Rabbit mAb (1:1000; CST, 3195), Vimentin Rabbit mAb (1:1000; abcam, Ab92547), Cleaved Caspase-3 pAb (1:1000; CST, 9661), GLUD1 pAb (1:1000; Proteintech, 14299-1-AP), GOT2 pAb (1:1000; Proteintech, 14800-1-AP), and α-Tubulin mAb (1:2000; Sigma, T5168).

### Reagents

The following reagents were used: N-Acetyl-L-cysteine (100 µM; Wako, 017-05131), Dimethyl 2-oxoglutarate (5 mM; Sigma, 349631), and 3-Mercaptopicolinic Acid (5 mM; Cayman chemical, 20895).

### Statistical analysis

Data are presented as the mean ± SEM. Statistical significance of the difference between experimental groups was assessed using an unpaired two-tailed Student’s t-test using GraphPad Prism 9 software. P values of < 0.05 were considered significant.

## Results

### The Expression of Both Variants of NFYA is Upregulated in Response to the Induction of Gluconeogenesis

Emerging evidence suggests that NFYA regulates glucose metabolism in the liver by regulating the expression of gluconeogenic enzymes (Zhang et al., 2018). Furthermore, several studies have reported that the gluconeogenic enzyme PCK1 is suppressed in cancer tissues and functions as a tumor suppressor in gluconeogenic organs such as the liver and kidney (Ma et al., 2013; Liu et al., 2018b; Tang et al., 2018). We compared the expression of the gluconeogenic enzymes PCK1 and G6PC with survival rates in several cancer types using the Kaplan-Meier plotter (Lánczky and Győrffy, 2021). The results showed a correlation between the expression of PCK1 and G6PC and prognosis in patients with HCC and clear cell renal cell carcinoma (ccRCC). In these cancer types, patients with high expression of PCK1 or G6PC showed a better prognosis than those with low expression (Supplementary Figures S1A, B). On the other hand, other cancer types did not show an obvious correlation (Supplementary Figures S1C, D). Although these findings led us to hypothesize that NFYA could play an essential role in tumor suppression of HCC through the regulation of gluconeogenesis, the regulatory mechanism for gluconeogenesis by NFYA in HCC and the functional differences between NFYA splicing variants remain unclear. To investigate the hypothesis, we first examined the expression of both variants of NFYA in the induction of gluconeogenesis. In the liver of C57BL/6N male mice fasted for 24 hours, the expression of both variants increased and decreased upon re-feeding (Figures 1A, B). This expression pattern was similar to that of the gluconeogenic enzymes PCK1 and G6PC, and the variation in the mouse blood glucose levels also indicated that the expression of the NFYA variants changes in response to gluconeogenesis (Figures 1A-C). These results suggest the involvement of NFYA in the regulation of gluconeogenesis in normal liver tissue, but differences between variants remain unclear.

**FIGURE 1.**
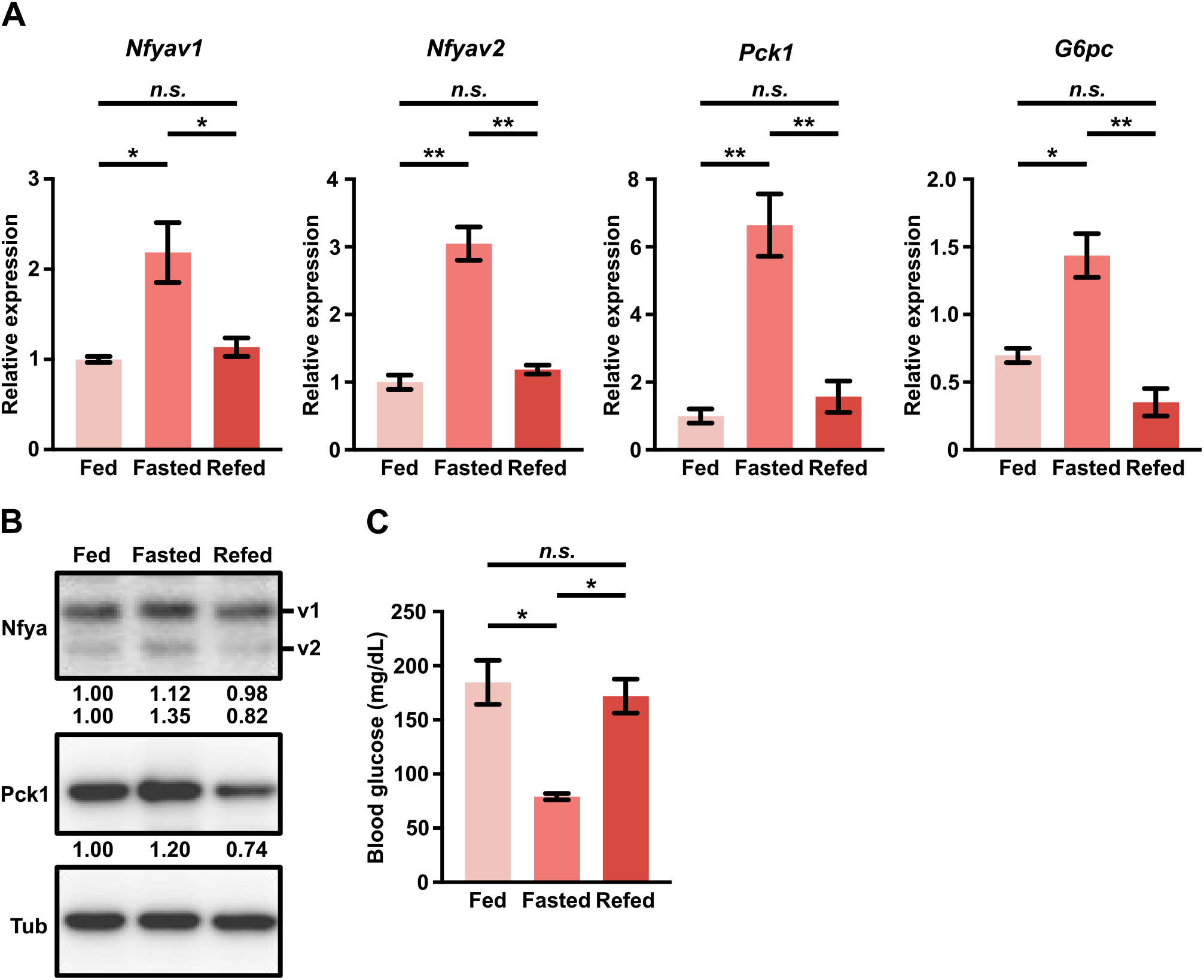
The expression of both variants of NFYA is upregulated in response to the induction of gluconeogenesis. **(A, B)** qRT-PCR (A; n=3) and western blot analysis (B) of Nfyav1, Nfyav2, Pck1, and G6pc in the liver from wild-type mice under fed, fasted, and re-fed conditions. The ratio of each band represents the relative amount of each protein normalized by Tubulin expression. **(C)** The blood glucose levels in wild-type mice under fed, fasted, and re-fed conditions (n=3). All error bars represent SEM; (n.s.) not significant; (*) P<0.05; (**) P<0.01.

### NFYAv2 Enhances the Anti-tumor Effects of Gluconeogenesis

To investigate the anti-tumor effects of gluconeogenesis regulation by NFYA on HCC, we examined the expression of NFYA and gluconeogenic enzymes in five different HCC cell lines. The expression of NFYA variants was associated with epithelial-mesenchymal transition (EMT) status as reported in breast cancer cells (Dolfini et al., 2019; Okada et al., 2022), but no association was observed for the expression of PCK1 and G6PC (Figure 2A). PCK1 and G6PC were highly expressed in HepG2 cells, moderately expressed in Hep3B, Huh4, and Huh-7 cells, and barely expressed in SK-Hep1 cells (Figure 2A). In subsequent experiments, SK-Hep1 cells, which do not express PCK1 and G6PC under normal conditions, were selected because we wanted to examine the anti-tumor effects of induced gluconeogenesis in HCC. Under glucose deprivation conditions, as expected, the anti-tumor effects of gluconeogenesis were confirmed by induction of the expression of PCK1 and cleaved caspase-3 (CC3) in SK-Hep1 cells (Figure 2B). Furthermore, given the higher induction of NFYAv2 compared to NFYAv1 under glucose deprivation conditions (Figure 2B), to investigate the functional differences among NFYA splicing variants, we generated NFYA-deficient and NFYAv1-specific deficient SK-Hep1 cells by CRISPR/Cas9 system (Figure 2C). We have reported that NFYAv1-specific deficient cells not only lack NFYAv1 expression but also overexpress NFYAv2 (Okada et al., 2022), which we will refer to as NFYAv2-overexpressing (NFYAv2OE) SK-Hep1 cells in subsequent experiments. Although NFYA-deficient cells inhibited cell proliferation under normal conditions, NFYAv2OE cells did not affect cell proliferation (Figure 2D). Using these cells, we examined the anti-tumor effects of induced gluconeogenesis under glucose deprivation conditions, and surprisingly, NFYAv2OE cells caused dramatic cell death. In addition, the induction of cell death was reduced in NFYA-deficient cells (Figures 2E-G). These observations suggest that NFYAv2, but not NFYAv1, is functionally important in the anti-tumor effects of gluconeogenesis induction.

**FIGURE 2.**
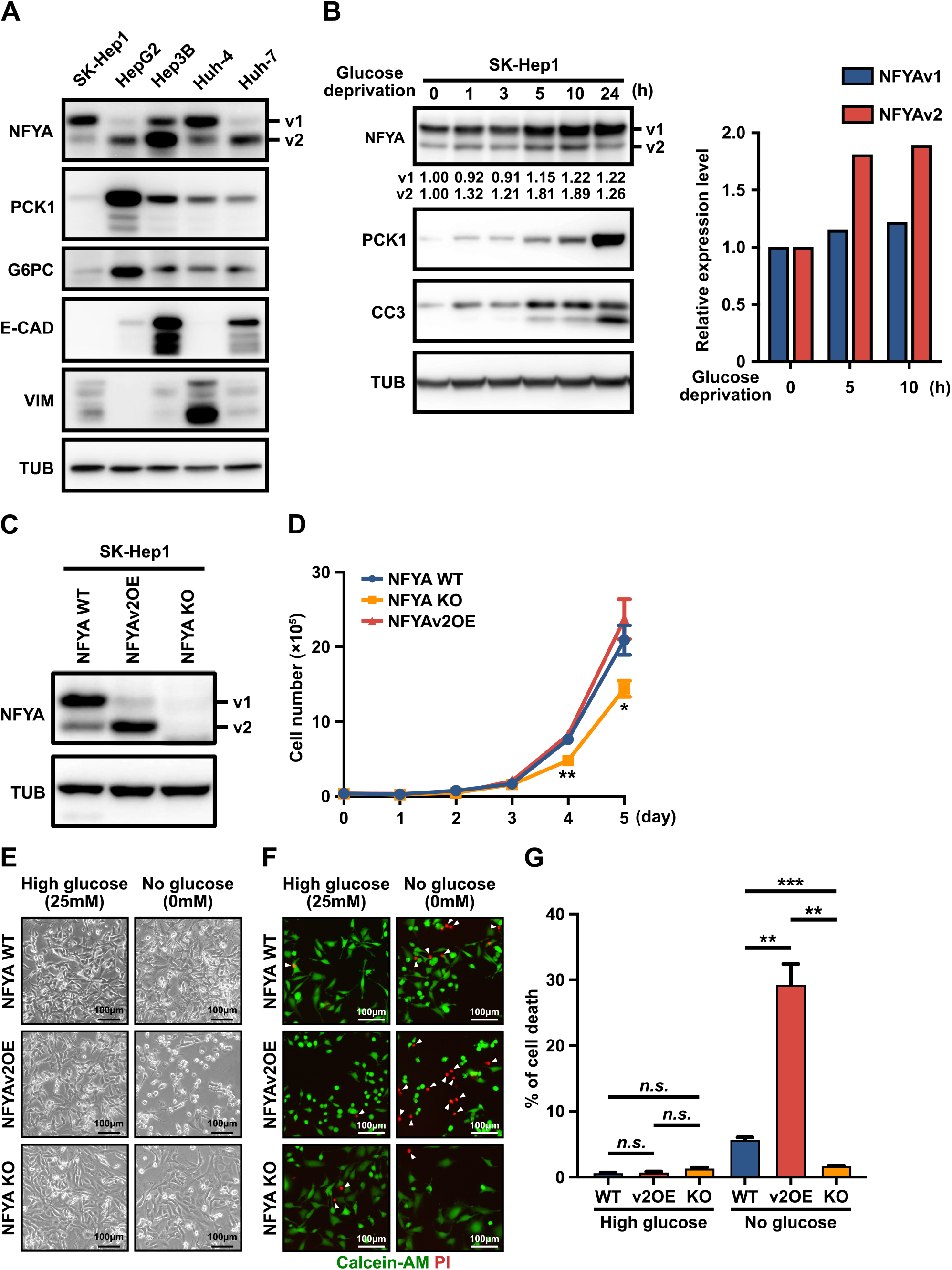
NFYAv2 enhances the anti-tumor effects of gluconeogenesis. **(A)** Western blot analysis of NFYA, gluconeogenic enzymes (PCK1 and G6PC), epithelial marker (E-CAD), and mesenchymal marker (VIM) protein levels in five HCC cell lines. **(B)** SK-Hep1 cells were cultured in glucose deprivation conditions for the indicated time. Western blot analysis of NFYA, PCK1, and CC3 protein levels in the cells. The intensity of each band represents the relative amount of NFYAv1 or v2 protein normalized by Tubulin expression. The bar graph represents the intensity of each band relative to the amount of Tubulin at 0, 5, and 10 hours of glucose deprivation. **(C)** Western blot analysis to validate the deletion of NFYA gene expression and NFYAv2 overexpression generated by CRISPR/Cas9 in SK-Hep1 cells. **(D)** The cumulative population of cells was measured for 5 consecutive days in wild-type, NFYAv2OE, and NFYA-deficient SK-Hep1 cells under normal conditions. **(E)** Representative images of cell morphology of wild-type, NFYAv2OE, and NFYA-deficient SK-Hep1 cells under normal or glucose deprivation conditions after 10 hours of culture. **(F)** Representative fluorescence images of living cells (green) detected with Calcein-AM and dead cells (red) detected with Propidium iodide (PI) in wild-type, NFYAv2OE, and NFYA-deficient SK-Hep1 cells under normal or glucose deprivation conditions after 10 hours culture. **(G)** The bar graph shows the percentage of dead cells among the more than 600 cells counted in each condition. All error bars represent SEM; (n.s.) not significant; (*) P<0.05; (**) P<0.01; (***) P<0.001.

### Anti-tumor Effects of Gluconeogenesis Enhanced by NFYAv2 are Caused by Oxidative Stress

Previous studies have reported that the anti-tumor effects of gluconeogenesis are achieved through oxidative stress induced by TCA cataplerosis (Liu et al., 2018b; Tang et al., 2018; Grasmann et al., 2019). Therefore, to reveal that the cell death enhanced by NFYAv2 is also caused by oxidative stress, we treated the cells with N-Acetyl-L-cysteine (NAC), an inhibitor of ROS. Treatment of wild-type, NFYA-deficient, and NFYAv2OE cells with NAC significantly blocked cell death under glucose deprivation conditions in all three cells (Figures 3A-C). Moreover, the addition of dimethyl α-ketoglutarate (α-KG), an intermediate in the TCA cycle, also significantly blocked cell death (Figures 3A-C). Indeed, NFYAv2OE cells increased ROS levels in glucose deprivation conditions, which was entirely abolished by NAC and α-KG (Figures 3D, E). These results indicate that NFYAv2 exerts the anti-tumor effect by promoting TCA cataplerosis under induced gluconeogenesis and causing cell death due to oxidative stress.

**FIGURE 3.**
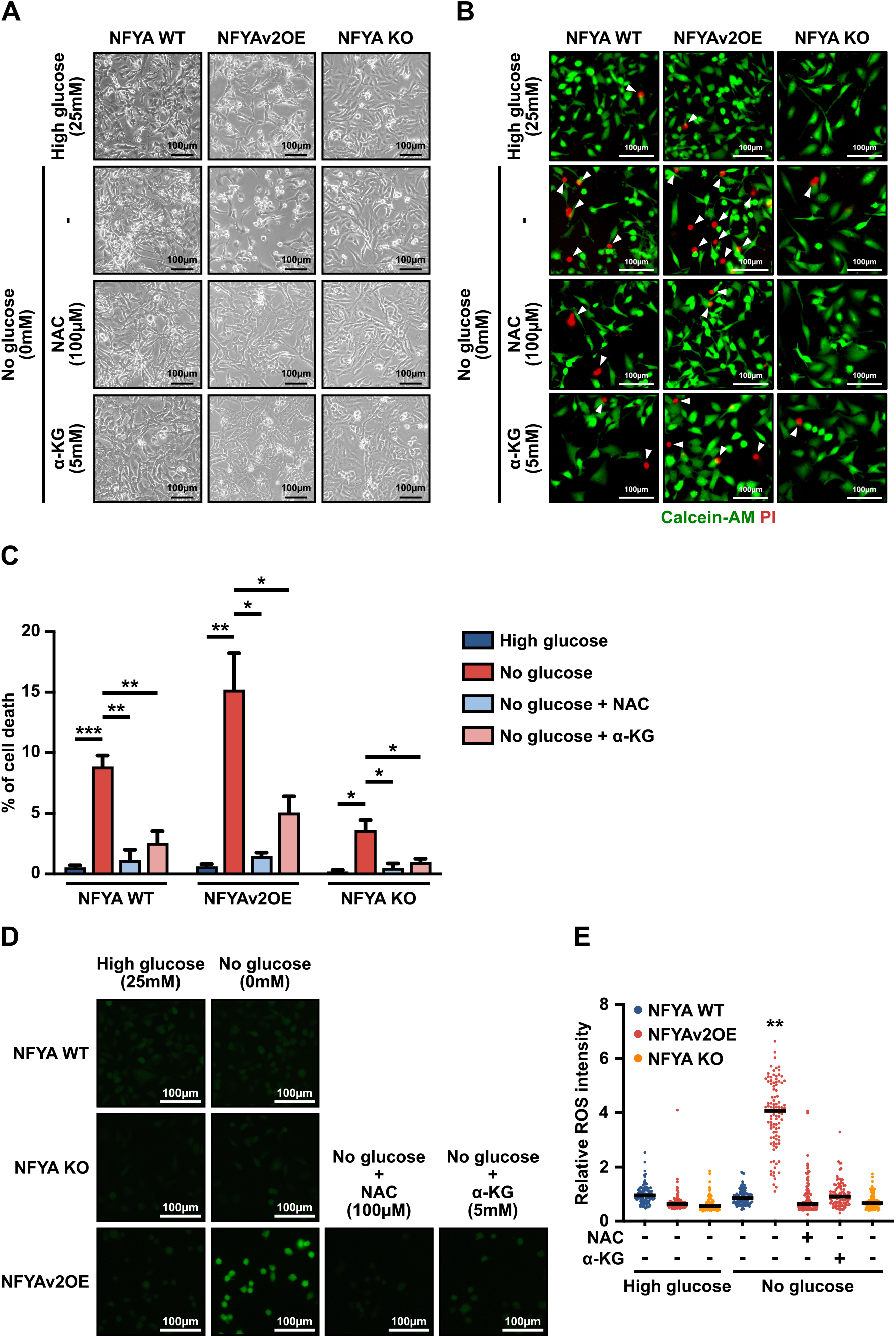
Anti-tumor effects of gluconeogenesis enhanced by NFYAv2 are caused by oxidative stress. **(A)** Representative images of cell morphology of wild-type, NFYAv2OE, and NFYA-deficient SK-Hep1 cells under normal or glucose deprivation conditions with treatment with NAC (100 µM) or dimethyl α-KG (5 mM) after 10 hours of culture. **(B)** Representative fluorescence images of living cells (green) detected with Calcein-AM and dead cells (red) detected with Propidium iodide (PI) in wild-type, NFYAv2OE, and NFYA-deficient SK-Hep1 cells under normal or glucose deprivation conditions with treatment with NAC (100 µM) or dimethyl α-KG (5 mM) after 10 hours culture. **(C)** The bar graph shows the percentage of dead cells among the more than 600 cells counted in each condition. **(D)** Representative fluorescence images of ROS (green) detected with highly sensitive DCFH-DA in wild-type, NFYAV2OE, and NFYA-deficient SK-Hep1 cells under normal or glucose deprivation conditions with treatment with NAC (100 µM) or dimethyl α-KG (5 mM) after 3.5 hours culture. **(E)** The dot plot shows the ROS intensity per cell among the 100 cells counted in each condition. Means represent SEM. All error bars represent SEM; (*) P<0.05; (**) P<0.01; (***) P<0.001.

### Glutaminolysis is Normal in NFYAv2OE SK-Hep1 Cells

Cancer cells rely on elevated glutaminolysis to maintain a functional TCA cycle (Wise and Thompson, 2010; Jin et al., 2016; Wang and Dong, 2019). Glutamine is intracellularly transported through the transporter SLC1A5 and is converted to glutamate by Glutaminase (GLS). Glutamate is converted to α-KG by Glutamate dehydrogenase 1 (GLUD1) and Glutamic-oxaloacetic transaminase 2 (GOT2) and becomes a resource for the TCA cycle (Supplementary Figure S2A). Therefore, we have not yet eliminated the possibility that the promotion of TCA cataplerosis by NFYAv2 is due to reduced glutaminolysis, not induced gluconeogenesis. We evaluated the expression of enzymes that catalyze glutaminolysis and the amount of intracellular glutamine, glutamate, and α-KG to eliminate the possibility. As a result, there were no differences in the expression of GLS, GLUD1, and GOT2 under glucose deprivation conditions between wild-type and NFYAv2OE cells (Figure 4C, Supplementary Figure S2B). Although the expression of SLC1A5 was decreased in NFYAv2OE cells, the amount of intracellular glutamine was not affected (Supplementary Figure S2C). In addition, the amounts of intracellular glutamate and α-KG were increased in NFYAv2OE cells compared to wild-type cells (Supplementary Figures S2D, E). However, there was no difference in the expression of Oxoglutarate dehydrogenase (OGDH), the gene encoding α-KG dehydrogenase, which catalyzed the conversion of α-KG to succinyl-CoA during the TCA cycle (Supplementary Figure S2F), indicating that these accumulations are the results of another mechanism and that glutaminolysis is generally functioning in NFYAV2OE cells. Therefore, the promotion of TCA cataplerosis by NFYAv2 may be due to the induction of gluconeogenesis.

**FIGURE 4.**
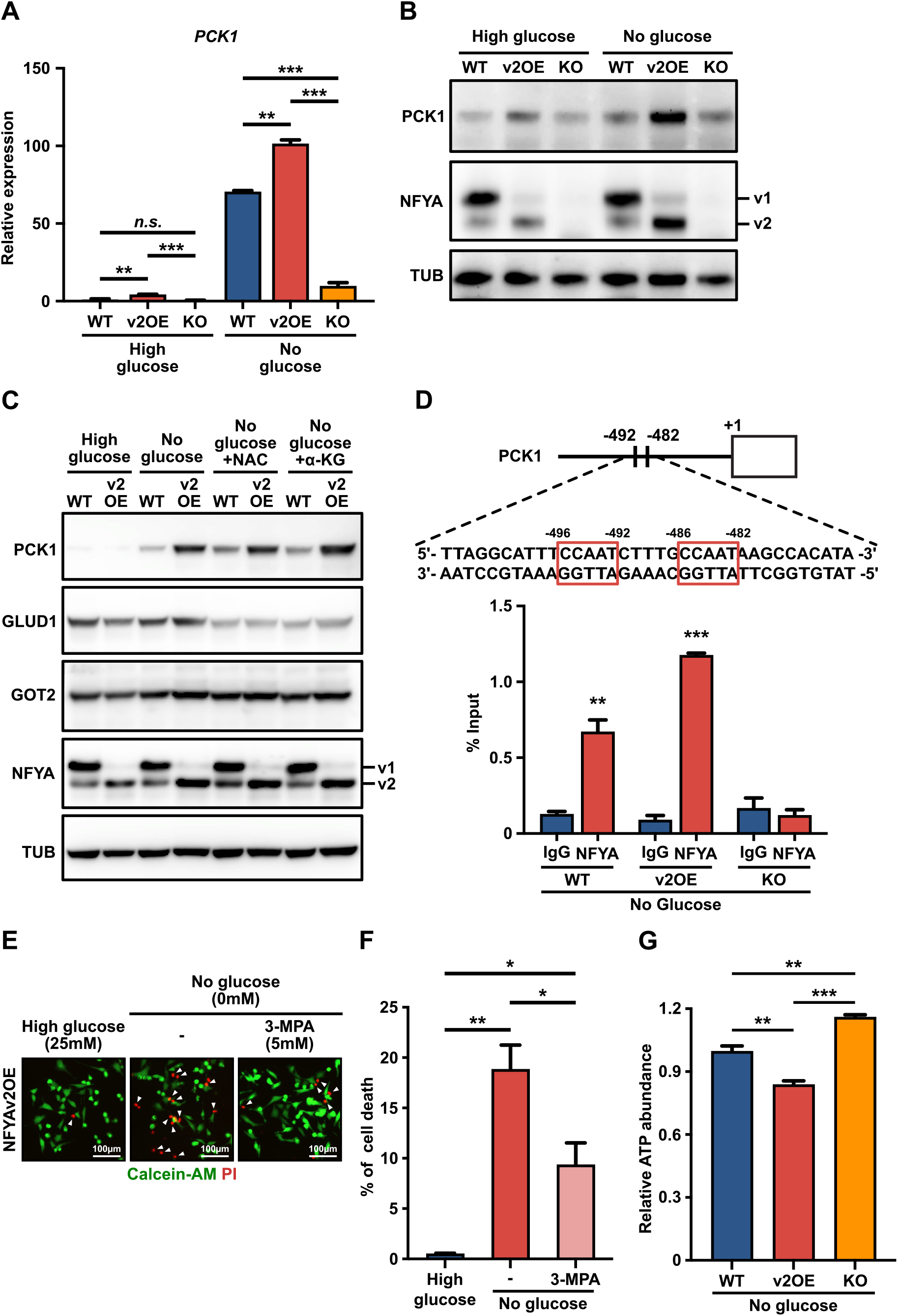
NFYAv2 enhances gluconeogenesis by transcriptional activation of PCK1. **(A, B)** qRT-PCR (A) and western blot analysis (B) of the expression levels of PCK1 in wild-type, NFYAv2OE, and NFYA-deficient SK-Hep1 cells under normal or glucose deprivation conditions after 10 hours of culture. **(C)** Western blot analysis of the expression levels of PCK1, GLUD1, GOT2, and NFYA in wild-type and NFYAv2OE SK-Hep1 cells under normal or glucose deprivation conditions with treatment with NAC (100 µM) or dimethyl α-KG (5 mM) after 10 hours of culture. **(D)** The location of NFYA binding sites on the PCK1 promoter region (upper panel). CUT&RUN assay to evaluate the association of NFYA with the promoter of PCK1 in wild-type, NFYAv2OE, and NFYA-deficient SK-Hep1 cells under glucose deprivation conditions after 5 hours of culture. Input and eluted chromatins were subjected to qRT-PCR analysis using promoter-specific primer. Data represents the % input of the immunoprecipitated chromatin for each gene (lower panel). **(E)** Representative fluorescence images of living cells (green) detected with Calcein-AM and dead cells (red) detected with Propidium iodide (PI) in NFYAv2OE SK-Hep1 cells under normal or glucose deprivation conditions with treatment with 3-MPA (5 mM) after 10 hours of culture. **(F)** The bar graph shows the percentage of dead cells among the more than 600 cells counted in each condition. **(G)** Intracellular ATP level in wild-type, NFYAv2OE, and NFYA-deficient SK-Hep1 cells under glucose deprivation conditions after 5 hours of culture. All error bars represent SEM; (n.s.) not significant; (**) P<0.01; (***) P<0.001.

### NFYAv2 Enhances Gluconeogenesis by Transcriptional Activation of PCK1

Given that the anti-tumor effects of NFYAv2 are achieved through gluconeogenesis and that NFYA regulates gluconeogenic enzyme PCK1 expression in hepatocytes (Zhang et al., 2018), we next examined the expression of PCK1 to determine how the NFYAv2 regulates gluconeogenesis in HCC. In NFYAv2OE cells, the induction of PCK1 expression caused by glucose deprivation treatment was significantly enhanced compared to wild-type cells (Figures 4A, B). Furthermore, the induction of PCK1 expression was significantly suppressed in NFYA-deficient cells (Figures 4A, B). The induction of PCK1 expression by NFYAv2 was not affected by treatment with NAC and α-KG, indicating that these inhibitors do not inhibit the induction of PCK1 expression (Figure 4C). These results indicate that NFYAv2 contributes to anti-tumor effects by controlling gluconeogenesis by regulating PCK1 expression. We further examined whether NFYA exerts the transcriptional regulation of PCK1 through direct binding to the promoter region by CUT&RUN assay using immunoprecipitation validated antibody. Since NFYA has been reported to bind to the CCAAT box in the promoter region of target genes (Dolfini et al., 2012), we searched for the CCAAT box in the promoter region of PCK1 and found it at 482 and 492 bp upstream (Figure 4D). Analysis of NFYA binding to these regions revealed that NFYA binds directly to the promoter region of PCK1 (Figure 4D). Interestingly, binding to the promoter region was higher in NFYAv2OE cells than in wild-type cells, supporting that NFYAv2 predominantly regulates PCK1 transcription, thereby contributing to its anti-tumor effects. In support of this notion, adding 3-Mercaptopicolinic acid (3-MPA), a PCK1 inhibitor (Balan et al., 2015; Brearley et al., 2020), significantly reduced cell death caused by glucose deprivation treatment (Figures 4E, F). Since glucose deprivation induces PCK1 expression in HCC, possibly causing stress due to energy depletion, we finally examined intracellular ATP levels. We found that ATP levels were significantly decreased in NFYAv2OE cells (Figure 4G).

## Discussion

It was reported that the expression of gluconeogenic enzymes PCK1, PCK2, and FBP1 predominantly decreased in HCC, and their forced expression induced apoptosis under glucose deprivation conditions, demonstrating the anti-tumor effect of gluconeogenesis in HCC (Yang et al., 2017; Liu et al., 2018b; Tang et al., 2018). Therefore, the gluconeogenesis pathway is a promising therapeutic target for HCC, which has a high mortality rate. However, there are still unsolved issues in the anti-tumor effects of gluconeogenesis induction.

First, the anti-tumor effects of gluconeogenesis are due to the forced expression of gluconeogenic enzymes, and the regulation of their expression is not well understood. Our analysis of NFYA splicing variants’ expression in HCC cells indicates NFYAv2 expression significantly increased under glucose deprivation conditions. Furthermore, we indicated that NFYAv2 induces high ROS levels in these cells, thereby promoting cell death under glucose deprivation conditions. It was also shown that this phenotype is due to NFYAv2 promoting transcription of a gluconeogenic enzyme, PCK1. In short, the NFYAv2-gluconeogenesis axis promotes cell death in HCC, suggesting that it may be helpful as a novel therapeutic target for HCC. However, the mechanism underlying the differential effects on target gene transcription among NFYA splicing variants remains elusive, and further studies are warranted.

The second issue is that the anti-tumor effects of gluconeogenesis enzymes are highly context-dependent. Previous studies have reported that gluconeogenesis has the anti-tumor effects in tumors derived from gluconeogenic organs, while it has the tumor-promoting effects in tumors derived from non-gluconeogenic organs (Grasmann et al., 2019; Wang and Dong, 2019). This study shows that NFYAv2 exerts its anti-tumor effect only under glucose deprivation conditions by regulating PCK1 transcription. Furthermore, we reported that NFYAv1 regulates the malignant behavior of breast cancer by regulating lipid metabolism using in vitro and in vivo models (Okada et al., 2022). Interestingly, we also have shown that NFYAv2 has no function in regulating breast cancer malignant behavior, and NFYAv2 has no tumor-promoting effects in tumors derived from the non-gluconeogenic organ (Okada et al., 2022). However, these effects may be due to the tissue-specific transcriptional regulation of target genes by NFYA; our study has not yet identified tissue-specific target genes of NFYA.

Finally, the most severe issue is safety as a therapeutic target. It has been reported that the expression of gluconeogenic enzymes is elevated in the liver of type 2 diabetes patients, resulting in increased gluconeogenesis (Yoon et al., 2001; Barthel and Schmoll, 2003; Sakai et al., 2012). Thus, forced expression of gluconeogenic enzymes increases the risk of type 2 diabetes and the anti-tumor effects on HCC, raising concerns about the safety as a therapeutic target. In this study, we show that the induction of PCK1 transcription by NFYAv2 is limited under glucose deprivation conditions. Since tumor tissues have extremely low glucose concentrations compared to normal tissues, likely, the transcriptional regulation of PCK1 by NFYAv2 occurs predominantly in tumor tissues. When Nfyav1-specific knockout mice, in which Nfyav2 overexpressed, were subjected to a pyruvate tolerance test, the behavior of blood glucose concentration was found to be unchanged compared to wild-type mice (Supplementary Figure S2G). These results suggest that NFYAv2 may be a valuable and attractive therapeutic target in therapy for HCC.

This study reveals a mechanism by which NFYAv2 promotes anti-tumor effects through the transcriptional regulation of PCK1 and induction of gluconeogenesis, indicating its importance as a therapeutic target for HCC.

## Data Availability Statement

The complete raw data sets supporting the findings of this study are available from the corresponding author upon reasonable request.

## Ethics Statement

The animal study was reviewed and approved by Okayama University Animal Care and Use Committee: OKU-2019325. Category/level: C (Experiments with mild stress or brief pain). Title of study: Functional analysis of Nfya in gluconeogenesis.

## Author Contributions

GT and NO conceived project and designed experiments. GT, RI, KY, CU, and NO performed experiments and analyzed data. NO procured funding and supervised the study. GT, RI, and NO wrote and edited the manuscript. All authors provided comments on the manuscript and gave final approval.

## Acknowledgments

We thank members of Division of Oncology and Molecular Biology, Kanazawa University and Medical Protein Engineering Laboratory, Okayama University for their help and input. Particularly, we thank Dr. Chiaki Takahashi for stimulating the discussion.

