## Supplementary information for "NFYA promotes the anti-tumor effects of gluconeogenesis in hepatocellular carcinoma through the regulation of PCK1 expression"

**Supplementary Table S1  Primer sequences for qRT-PCR analyses.**

| **qRT-PCR primer sequences** | |
|---|---|
| mouse Actb forward primer | 5'- GATCTGGCACCACACCTTCT -3' |
| mouse Actb reverse primer | 5'- GGGGTGTTGAAGGTCTCAAA -3' |
| mouse Nfyav1 forward primer | 5'- AAGTCCAGACCCTCCAGGTAGT -3' |
| mouse Nfyav1 reverse primer | 5'- GATGGGTTGGCCTGTTGAT -3' |
| mouse Nfyav2 forward primer | 5'- GCCATGGAGCAGTATACGACA -3' |
| mouse Nfyav2 reverse primer | 5'- CCTGGACCTGCTGCTGAA -3' |
| mouse Pck1 forward primer | 5'- CCTTTGGAAGCGGATATGGT -3' |
| mouse Pck1 reverse primer | 5'- TTGCCTTCGGGGTTAGTTATG -3' |
| mouse G6pc forward primer | 5'- ACTGTGGGCATCAATCTCCTCT -3' |
| mouse G6pc reverse primer | 5'- GGGCGTTGTCCAAACAGAA -3' |
| human ACTB forward primer | 5'- ACCAACTGGGACGACATGGAGAAA -3' |
| human ACTB reverse primer | 5'- TAGCACAGCCTGGATAGCAACGTA -3' |
| human PCK1 forward primer | 5'- GACATTGCCTGGATGAAGTTTG -3' |
| human PCK1 reverse primer | 5'- TTCTTCTGGATGGTCTTGATGG -3' |
| human SLC1A5 forward primer | 5'- TCCGCTTCTTCAACTCCTTCA -3' |
| human SLC1A5 reverse primer | 5'- AAACCCACATCCTCCATCTCC -3' |
| human GLS forward primer | 5'- TTCTCAGGGCAGTTTGCTTTC -3' |
| human GLS reverse primer | 5'- TTGCCCATCTTATCCAGAGGA -3' |
| human GLUD1 forward primer | 5'- AAGGCAAAGCCCTATGAAGGA -3' |
| human GLUD1 reverse primer | 5'- CATTGGCACCTTCAGCAATG -3' |
| human GOT2 forward primer | 5'- ACCGGGATGATAATGGAAAGC -3' |
| human GOT2 reverse primer | 5'- CTTGCAAAATTCAGCCAGTCC -3' |
| human OGDH forward primer | 5'- GCAGATGTGCAACGATGACC -3' |
| human OGDH reverse primer | 5'- TGGAAGAAGTTGCCAGGAGTG -3' |
| human PCK1 forward primer for CUT&RUN assay | 5'- GGTGCATCCTTCCCATGAAC -3' |
| human PCK1 reverse primer for CUT&RUN assay | 5'- CTGGTTGGCAAAACACCACA -3' |

### Tsujimoto and Ito *et al.* Supplementary Figure S1

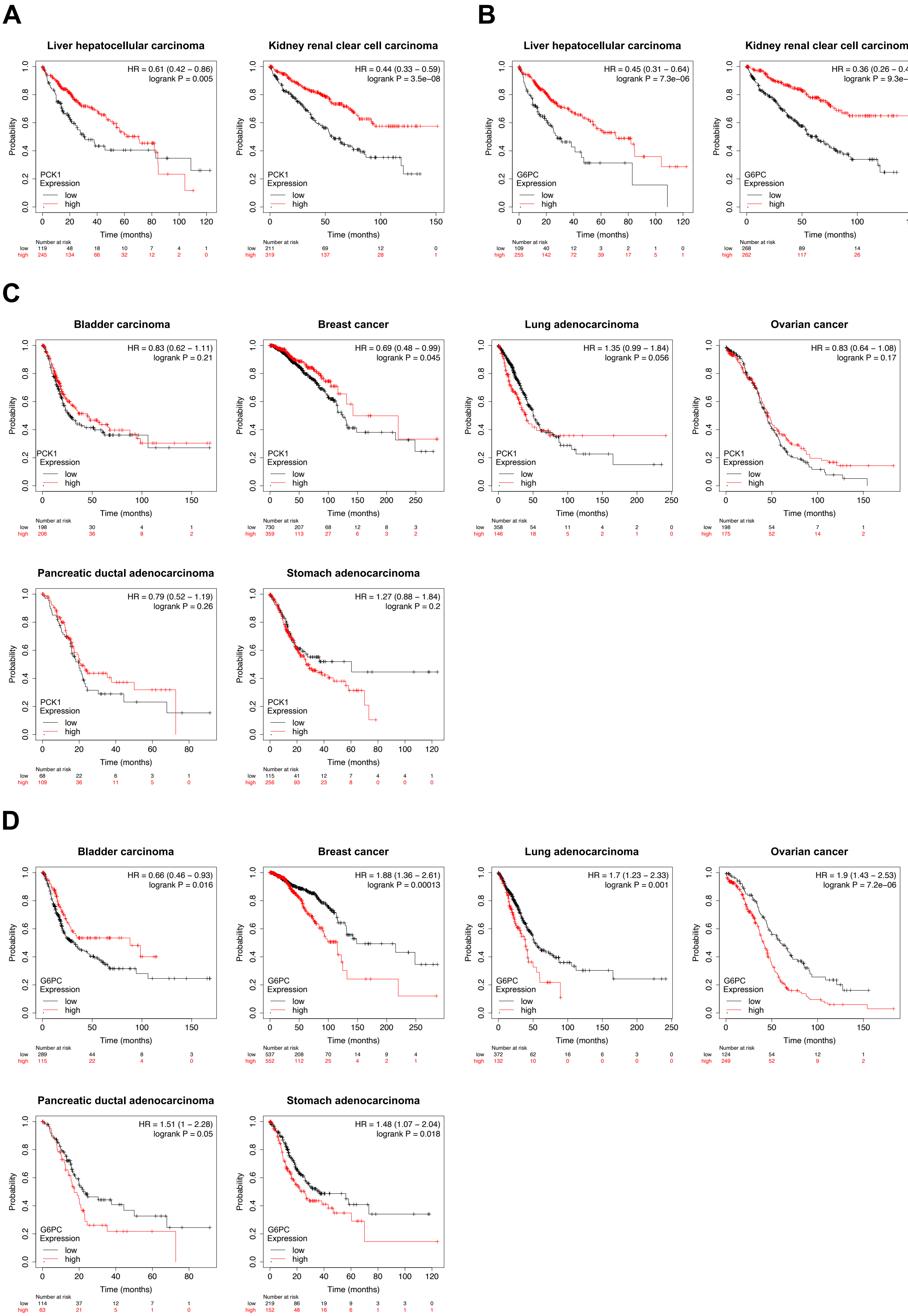

**Supplementary Figure S1.** The expression of PCK1 and G6PC predicts better patient survival only for gluconeogenic organ cancer. Kaplan-Meier plots of overall survival of patients with gluconeogenic organ cancer (A; for PCK1, B; for G6PC) and non-gluconeogenic organ cancer (C; for PCK1, D; for G6PC). Data were obtained from the Kaplan-Meier plotter online tool. The hazard ratio (HR) and respective log-rank p-values identifying the high-expression group (red) and the low-expression group (black) are shown.

### Tsujimoto and Ito *et al.* Supplementary Figure S2

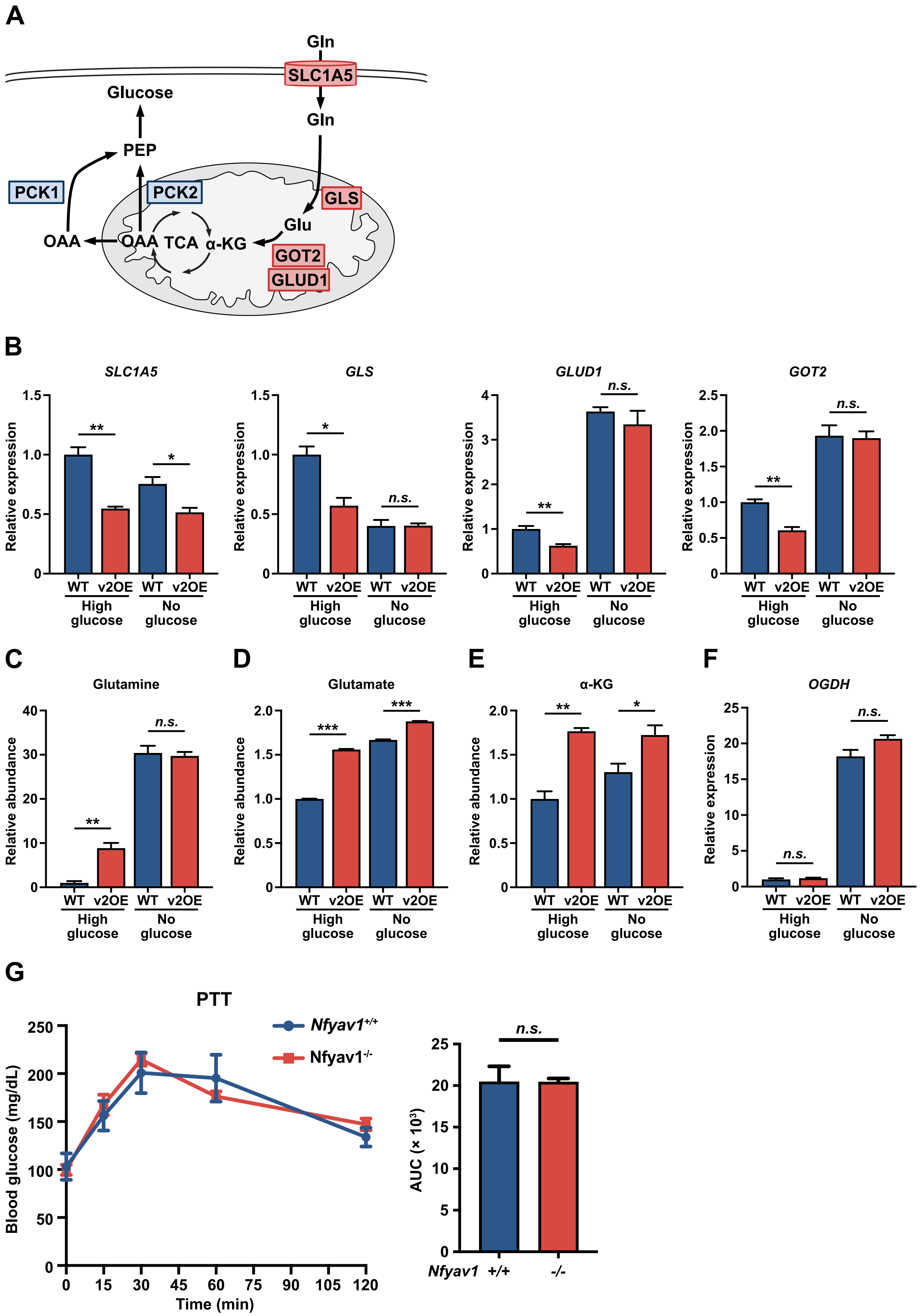

**Supplementary Figure S2.** Glutaminolysis is normal in NFYAv2OE SK-Hep1 cells. **(A)** A diagram illustrating the reaction of glutaminolysis and gluconeogenesis. **(B)** qRT-PCR analysis of the expression levels of SLC1A5, GLS, GLUD1, and GOT2 in wild-type and NFYAv2OE SK-Hep1 cells under normal or glucose deprivation conditions after 5 hours of culture. **(C-E)** The intracellular glutamine (C), glutamate (D), and a-KG (E) levels in wild-type and NFYAv2OE SK-Hep1 cells under normal or glucose deprivation conditions after 5 hours of culture. **(F)** qRT-PCR analysis of the expression levels of OGDH in wild-type and NFYAv2OE SK-Hep1 cells under normal or glucose deprivation conditions after 5 hours of culture. **(G)** Pyruvate tolerance test (PTT) in *Nfyav1*$^{+/+}$ (n=5) and *Nfyav1*$^{-/-}$ (n=4) mice. A bar graph shows the area under curve. All error bars represent SEM; (n.s.) not significant; (*) P<0.05; (**) P<0.01; (***) P<0.001.
